# Narcosis biosensor for the detection of bacterial membrane disruption by naphthenic acids

**DOI:** 10.64898/2026.05.22.727335

**Authors:** Tyson Bookout, Shawn Lewenza

## Abstract

Naphthenic acids are amphipathic compounds whose toxicity has primarily been attributed to narcosis toxicity to cell membranes. However, few methods exist that specifically study the membrane disruption and toxicity of this complex family of cyclic, polycyclic and acyclic alkyl-substituted carboxylic acids. Here we describe a whole cell biosensor approach that relies on the ability of *Pseudomonas aeruginosa*, a ubiquitous environmental organism and opportunistic pathogen, to sense membrane damage (narcosis) and induce protective genes to repair and protect the outer membrane. Many classes of membrane disrupting antimicrobials induce the expression of two operons that encode protective defense systems against outer membrane (OM) damage, including antimicrobial peptides, chelators, and detergents. We demonstrate that the *pmrF* and *spdE2* transcriptional *lux* reporters are induced by exposure to individual NA compounds with diverse structures, as well as mixtures and naphthenic acid fraction compounds (NAFCs). To further support the narcosis hypothesis, we demonstrated that NA permeabilizes the outer membrane to assist in lysozyme killing, and disrupts the inner membrane integrity, allowing uptake of the DNA binding dye propidium iodide. The conventional OM permeability assay that measures NPN fluorescence is not applicable to study NAs, because they stimulate NPN fluorescence in the absence of cells. This narcosis biosensor approach constitutes a rapid and simple method to measure narcosis and could be developed as a novel toxicity indicator of oil sands tailings.

## Introduction

Naphthenic acids (NAs) are a complex mixture of cyclic, polycyclic and acyclic alkyl-substituted carboxylic acids that accumulate in oil sands tailings after the extraction of bitumen from the oil sands using the Clark process of alkaline water extraction (1). Naphthenic acids can accumulate in oil sands process-affected water (OSPW) between 20-100 mg/L and their toxicity has been assessed in diverse animal models that include amphibians, fish, birds, mammals, and water fleas (2–5). The primary mechanism of toxicity of naphthenic acids is attributed to narcosis, due to their ability to integrate into and disrupt cell membranes (6–8, 5, 9). NA are amphipathic compounds due to their charged carboxylic acids and possess hydrophobic cycloalkane ring structures and alkyl groups, which result in membrane disrupting, surfactant properties (6).

However, very few studies use specific methods to assess membrane damage. The luminescent, bacterium *Vibrio fischeri* (Microtox) is used as a general toxicity indicator of OSPW and naphthenic acids, where the loss of luminescence indicates toxicity. Several Microtox studies attributed the increased toxicity of larger molecular weight fatty acid and single ring NA compounds to an increased narcosis effect (3), since increased molecular weight is consistent with increased hydrophobicity and toxicity. Another bacterial study demonstrated that *Rhodococcus aertherivorans* exposed to naphthenic acids showed slower growth, morphological changes, produced altered membrane fatty acids and took up the impermeable, DNA binding dye propidium iodide (PI), all of which indicated membrane damage effects (10). A microscopy study on the toxicity of naphthenic acids to plant cells provided several lines of evidence that NA were acting on plant cell and organelle membranes (9). Naphthenic acid exposure to plants dramatically altered the shape and dynamics of several membrane-bound organelles, evidence of NA partitioning into and disrupting membranes (9).

The outer bacterial membrane is the first barrier of entry to antimicrobials, and there are numerous mechanisms of reducing the outer membrane permeability to antibiotics (11). One resistance mechanism in *P. aeruginosa* is the modification of lipopolysaccharide (LPS) in the outer leaflet of the outer membrane to limit the permeability and entry of antimicrobial peptides; components of the innate immune systems of insects, vertebrates, and plants (12, 13). The addition of aminoarabinose (L-Ara4N) to lipid A phosphates promotes antimicrobial resistance by neutralizing negative surface charges, reducing the binding and membrane disruption by cationic antimicrobial peptides and chelators (14, 15, 12, 16). *P. aeruginosa* also produces the polycation spermidine on the cell surface which masks negative surface charges, stabilizes LPS as a substitute for divalent cations, and contributes to antimicrobial peptide resistance (15, 17, 18).

We recently used transcriptional reporters to the *pmr* operon (*PA3552-PA3559*; *arn*) and *spd* operon (*PA4773-PA4774; speD2E2*) that encode the L-Ara4N modification and spermidine production, respectively, as biosensors for detecting generic outer membrane damage and for the discovery of novel, membrane-acting antimicrobials (19). The *pmr* and *spd* operons were strongly induced during exposure to diverse outer membrane damaging compounds that included antimicrobial peptides, chelators, and detergents (19). The biosensor approach was proposed as a stand-alone, alternative, simple and high throughput method to the NPN uptake assay, a standard method used to measure OM damage (19). Here, we report on the use of *pmr* and *spd* transcriptional fusions as a biosensor approach to identify naphthenic acids as direct, narcosis causing compounds.

## Methods and Materials

### Strains and biosensor gene expression assays

The *pmrF::lux* and *speE2::lux* reporters are mini-Tn*5*-*lux* transposon insertion mutants and transcriptional reporters that were previously constructed (15) and used extensively as indicators of outer membrane damage by antimicrobials (14, 15, 20, 16, 19). Overnight biosensor cultures were grown in LB media + 50 µg/mL tetracycline at 37°C and sub-cultured into 10 mL of dilute LB (1/32 v/v). Stock solution of model NA compounds were prepared in solutions of 55% polyethylene glycol 400 and 45% ethanol (PEGEt) as previously described (21). In black 96-well, clear bottom plates (Thermo Scientific), NA were diluted to 125 µg/mL in 99 µl Basal Minimal Medium (BM2) (0.5 mM Mg^2+^) with 20 mM succinate (22) and inoculated with 1 µL of the biosensor culture. To measure the biosensor response to the NAFC extracts (prepared in 50% acetonitrile + 0.1% NH_4_OH and chemically analyzed by high resolution Orbitrap mass spectrometry) (23, 24), a 10 µL aliquot of an extract was dispensed into 90 µL BM2 media using the same inoculum in black 96-well microplates. A Breath-Easy® (Sigma-Aldrich) membrane was used to seal the microplate to lower evaporation rates during a 15-hour protocol in a PerkinElmer 1420 Multilabel Counter Victor^3^ (2 sec shake; gene expression in counts per second (CPS), and growth (OD_600_), all measured 45 times every 20 minutes). The biosensor gene expression was normalized to growth (CPS/OD_600_) for each read, and Fold Gene Expression was calculated by dividing the CPS/OD_600_ values of the NA treated sample by those of the untreated control (containing 0.625% PEGEt or 5% acetonitrile). One-way ANOVA and Dunnett T-tests were used to calculate statistical significance compared to the untreated sample.

### Biosensor spot agar plate

Biosensor cultures were grown overnight in LB media + 50 µg/mL tetracycline and diluted in LB to an OD_600_ of 0.5. Sterile cotton swabs were used to spread a bacterial lawn onto diluted LB (1/32) agar plates. Diluted LB was used to mimic nutrient limited growth medium for optimal NA detection. Aliquots of 4 µL of a 20 mg/mL stock of naphthenic acid model compounds, mixtures or additional hydrocarbons were spotted in a grid on top of the bacterial lawn. The petri dish was incubated for 21 hours at 37°C and imaged using a ChemiDoc (Bio-Rad) to visualise bioluminescent patterns surrounding each spot.

### Outer membrane damage measured by NPN uptake

1-N-phenylnaphylamine (NPN) is a fluorescent dye when integrated into the hydrophobic environment of disturbed bacterial membranes (25, 19). Mid-log cultures of PAO1 (grown in LB at 37°C) were centrifuged at 8000 rpm for 3 minutes, and pellets were resuspended in equal volumes of a buffer containing 5mM HEPES (pH 7.2), 5mM glucose, and 0.2% sodium azide for disabling active efflux. NPN was added at a final concentration of 0.01 mM, followed by an addition of hydrocarbon compounds for a final concentration of 125 mg/L. Green fluorescence was measured in a Spectra Max M2 spectrophotometer at 5 second intervals for 60 seconds, using the excitation and emission wavelengths of 350 nm and 420 nm, respectively. NPN (0.01 mM) was also added to the buffer containing 125 µg/mL hydrocarbons to measure baseline fluorescence.

### Inner and outer membrane damage measured by PI uptake

PAO1 cultures were grown to mid-log phase in LB at 37°C. Cells were pelleted (8000 rpm for 3 minutes) and washed once in equal volumes of phosphate buffered saline (PBS). Propidium iodide (PI) was added to a final concentration of 5 µg/mL. Naphthenic acids (at a final concentration of 125 µg/ml) were mixed with aliquots of 100 µL bacterial cells in a black, clear bottom 96-well microplate. Using a PerkinElmer 1420 multilabel counter Victor^3^, red fluorescence was measured every 20 minutes for 15 hours (using excitation and emission wavelengths of 555 and 632 nm, respectively) as an indicator of PI uptake into the cell after disruption of both outer and inner membranes. Bacteria can take up PI but remain viable, so it is not strictly an indicator of non-viable cells (26).

### Lysozyme assisted bacterial killing

Using minor modifications of a previously described method (27), *P. aeruginosa* PAO1 cultures (3 mL) were grown to mid-log phase (~3.5 hours in LB at 37°C) and normalized to an OD_600_ of 0.4. Cells were pelleted by centrifugation (8000 rpm for 3 minutes) and washed twice in 3 mL (PBS). Aliquots of 80 µL were dispensed to each well of a clear, 96-well microplate, and mixed with 10 µL of a naphthenic acid (NA) compound for each concentration indicated and lysozyme solution (final 1 mg/ml), and cell lysis was measured by monitoring OD_600_ every 5 min for 75 minutes in the Victor^3^ 1420 Multilabel Counter (PerkinElmer). Control reactions containing only lysozyme, or only the OM permeabilizing agents did not cause cell lysis.

## Results

### Naphthenic acids induce expression of the *pmrF* and *spdE2* transcriptional reporters of protective modifications to the outer membrane

*P. aeruginosa* senses outer membrane damage caused by diverse antimicrobials that target the outer membrane and induces the expression of multiple surface modifications that function to repair and protect the membrane (14, 15, 20, 16, 19). Naphthenic acids are amphipathic compounds with charged and hydrophobic features, which are proposed to act by damaging the cell membrane through a narcosis mechanism (6–8, 5, 9). Here we wanted to determine if the narcosis by naphthenic acids was detectable using the *pmrF* and *spdE2* transcriptional *lux* reporters, and to validate that naphthenic acids damage bacterial membranes.

We tested a large collection of model NA compounds, commercial and NAFC mixtures, and acyclic fatty acids for the ability to induce expression of the *pmrF::lux* transcriptional reporter (Fig 1). The model naphthenic acid compounds included mostly classic O_2_compounds with single rings, aromatic NAs, and an O_4_ NA compound (Table 1). When added to defined, liquid minimal media at 125 mg/L, most naphthenic acids compounds significantly induced *pmrF*::lux expression (Fig 1A). This NA concentration is sub-lethal, and below the concentration needed to inhibit growth. Related hydrocarbons that included BTEX, terpenes (branched alcohols) and alkanes were also tested, and only octane, decane and citronellol were shown to induce the outer membrane damage-responsive *pmrF* promoter. A panel of fatty acids did not induce *pmrF::lux* expression in liquid culture (Fig 1A).

**Table 1.** Summary of naphthenic acid model compounds and hydrocarbons used in this study.

| Group | Compound | Acronym | Description |
| --- | --- | --- | --- |
| **NAs** | Cyclopentane carboxylic acid | CPCA | classic O<sub>2</sub> compound |
|  | Cyclohexane pentanoic acid | CHPA | classic O<sub>2</sub> compound |
|  | Cyclohexane acetic acid | CHAA | classic O<sub>2</sub> compound |
|  | 1-Adamantanecarboxylic acid | ACA | classic O<sub>2</sub> compound |
|  | Cyclohexane carboxylic acid | CHCA | classic O<sub>2</sub> compound |
| **Aromatic** | 3-Phenylpropionic acid | PPA | aromatic O<sub>2</sub> compound |
|  | 4-Dimethylphenylacetic acid | DMPAA | aromatic O<sub>2</sub> compound |
|  | 5,6,7,8-Tetrahydro-2-naphthoic acid | THNA | aromatic O<sub>2</sub> compound |
|  | Fusaric acid | FA | aromatic O<sub>2</sub> compound |
| **NAs** | Cyclohexane dicarboxylic acid | CHDCA | O<sub>4</sub> compound |
|  | Cyclohexyl succinic acid | CHSA | O<sub>4</sub> compound |
| **Fatty acids** | Hexanoic acid | HA/C6 | acyclic fatty acid O<sub>2</sub> |
|  | Citronellic acid | CA | acyclic fatty acid O<sub>2</sub> |
|  | Decanoic acid | DA | acyclic fatty acid O<sub>2</sub> |
|  | Dodecanoic acid | C12 | acyclic fatty acid O<sub>2</sub> |
|  | Pentadecanoic acid | C15 | acyclic fatty acid O<sub>2</sub> |
|  | Stearic acid | C18 | acyclic fatty acid O<sub>2</sub> |
|  | Oleic acid | C18:1 | acyclic fatty acid O<sub>2</sub> |
|  | Arachidic acid | C20 | acyclic fatty acid O<sub>2</sub> |
| **Alkanes** | Hexane | HEX | alkane |
|  | Octane | OCTA | alkane |
|  | Decane | DECA | alkane |
|  | Dodecane | DODE | alkane |
|  | Octadecane | OCT | alkane |
|  | Methylcyclohexane | MCM | cyclic alkane |
|  | Butylcyclohexane | BCM | cyclic alkane |
| **BTEX** | Benzene | B | aromatic |
|  | Toluene | T | aromatic |
|  | Ethylbenzene | E | aromatic |
|  | Xylene | X | aromatic |
| **Terpenes** | Citronellol | CIT | acyclic terpene |
|  | Geraniol | GER | acyclic terpene |
| **Mixes** | Acyclic NA mix (Sigma) | Sigma | Commercial NA mix with abundant FAs |
|  | Acyclic NA mix (Acros) | Acros | Commercial NA mix with abundant FAs |
|  | Custom NA mix | 9xMix | HA, DA, CPCA, CHPA, CHAA, ACA, PPA, DMPAA, THNA |
|  | OSPW | NAFC | SPE extract from OSPW sample |

**Figure 1.**
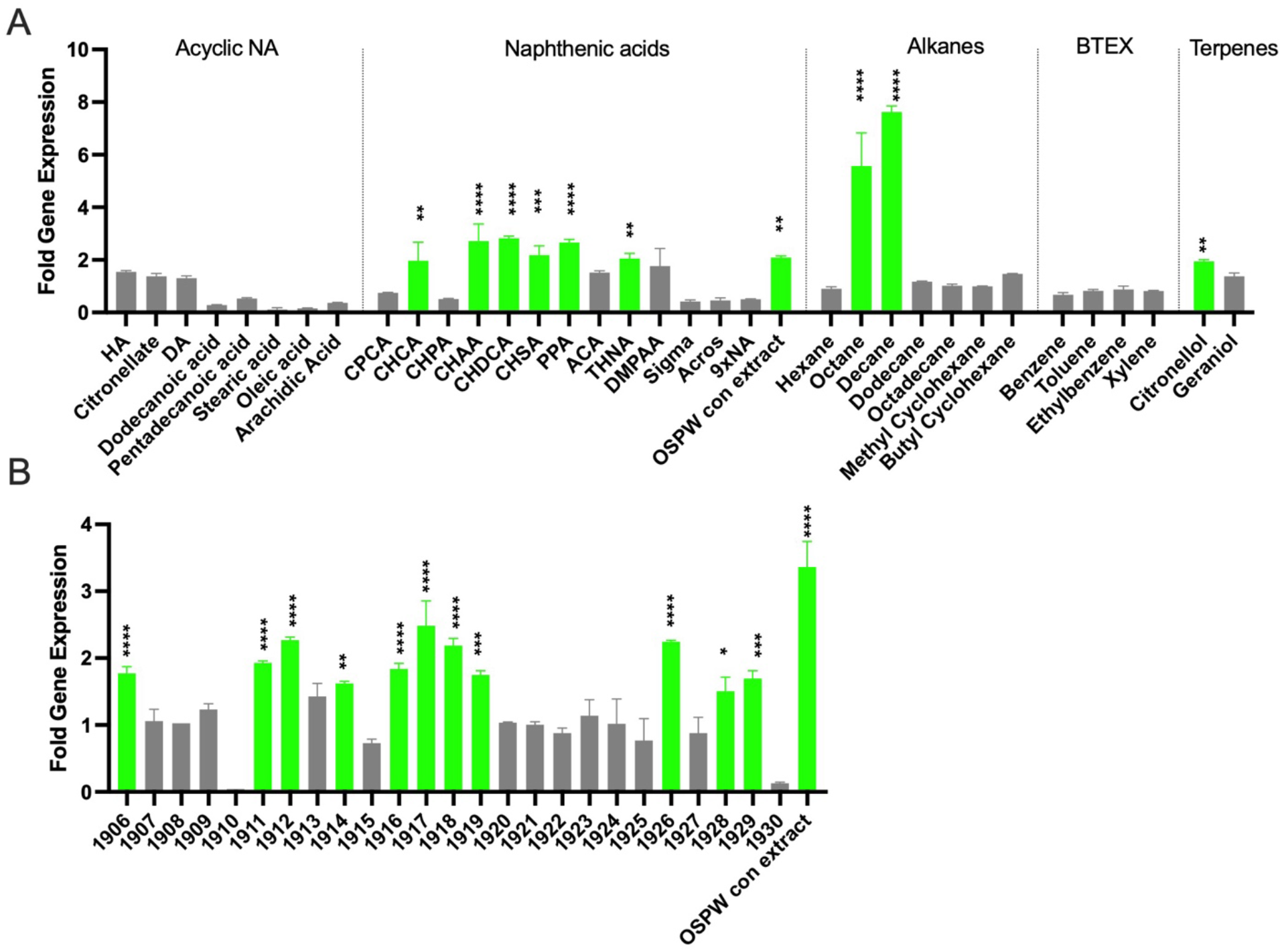
Narcosis biosensor response of the *pmrF::lux* reporter to naphthenic acids. **A)** Individual model NA compounds and related hydrocarbons were tested for biosensor detection by exposure to 125 mg/L of each compound. See Table 1 for full names. Various mixtures were also tested including “Sigma” and “Acros”, two commercially available NA mixtures, “9xNA” refers to a custom mixture of seven NAs and two fatty acids, and “OSPW con extract”, a reference NAFC extract from OSPW. The Fold Gene Expression is the maximum induction values of gene expression upon exposure, representing the average and standard deviation of triplicate values taken at 7 hours. **B)** A panel of 26 NAFC extracts from oil sands industry water samples were tested for biosensor detection. Each extract was diluted 1/10 into BM2 media. All values represent the average and standard deviation of triplicate values taken at 4 hours, and all experiments were repeated at least three times. Data were analyzed by one-way ANOVA to test for an overall treatment effect, followed by Dunnett’s multiple-comparisons test to compare each treatment with the control. Significant gene induction relative to the control is shown in bright green with asterisks (* p < 0.05, ** p < 0.01, *** p < 0.001, **** p < 0.0001).

We also tested NA mixtures, including multiple NAFC extracts derived from OSPW, two commercially available NA mixtures, and a custom mixture of seven NAs and two short-chain fatty acids. Only the solid phase extracts of OSPW induced expression of the *pmrF::lux* biosensor in liquid culture (Fig 1). Roughly half of the panel of OSPW extracts induced expression of the *pmrF::lux* biosensor, when tested in total NAFC concentrations ranging from 30-400 mg/L (Fig 1B). Induction of the narcosis biosensor to NAFC mixtures was not concentration dependent and therefore was likely dependent on the overall composition of NA compounds, requiring sufficient levels of membrane-damaging NA compounds.

The *pmrF::lux* biosensor has a wide dynamic range, which can distinguish weak and strong OM disrupters, and has dose-dependent responses to increasing concentrations of OM targeting compounds (19). Figure 2 demonstrates the dose response effects upon exposure to increasing concentrations of model NA compounds, which resulted in linear increases in gene expression output (Fig 2A). When the pH was adjusted to 6.5 after the addition of NA, the narcosis effects were strongest at the highest concentrations (Fig 2B). These results demonstrate that the *pmrF* biosensor can be used semi-quantitatively to report on the relative amount of outer membrane damage.

**Figure 2.**
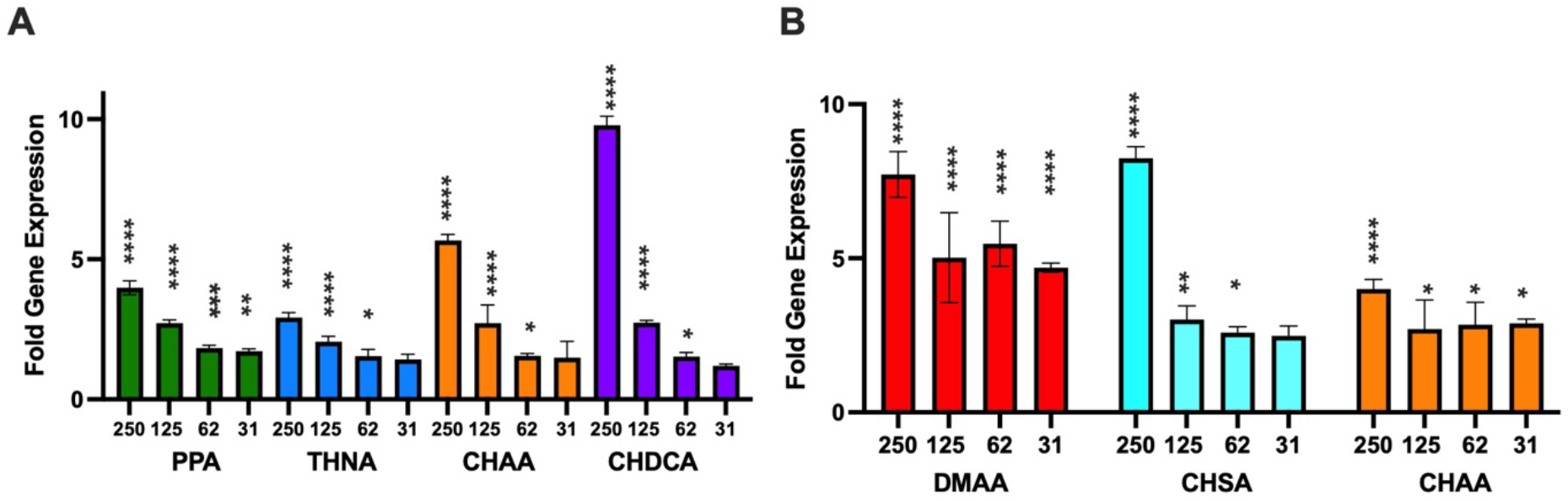
Dose responses of narcosis biosensor exposed to increasing concentrations of naphthenic acids. The *pmrF::lux* biosensor was exposed to individual NA model compounds at concentrations 31, 62, 125, and 250 mg/L in A) BM2 media and B) BM2 media that was pH adjusted to 6.5. See Table 1 for full names. All values represent the average and standard deviation of triplicate values taken at 4 hours, and all experiments were repeated at least three times. Data were analyzed by one-way ANOVA to test for an overall treatment effect, followed by Dunnett’s multiple-comparisons test to compare each treatment with the control. Significant gene induction relative to the control is shown with asterisks (* p < 0.05, ** p < 0.01, *** p < 0.001, **** p < 0.0001).

We next tested the addition of NA in a solid agar assay where the biosensor is spread as a lawn onto an agar surface, and plates were imaged after overnight growth for luminescence patterns at spot locations (Fig 3). Here we tested both *pmrF* and *speE2* reporters, which previously were shown to both respond similarly to a large panel of OM disrupting compounds (19). Both reporters responded to all model NA compounds (Fig 3A), and interestingly to all fatty acid compounds ranging in length from C6-C20 (Fig 3B). Antimicrobial activity is seen as a zone of killing in the centre of the spot and a ring of luminescence that surrounds the killing zone, which is a pattern consistent with biosensor detection at concentrations near the minimal inhibitory concentration.

**Figure 3.**
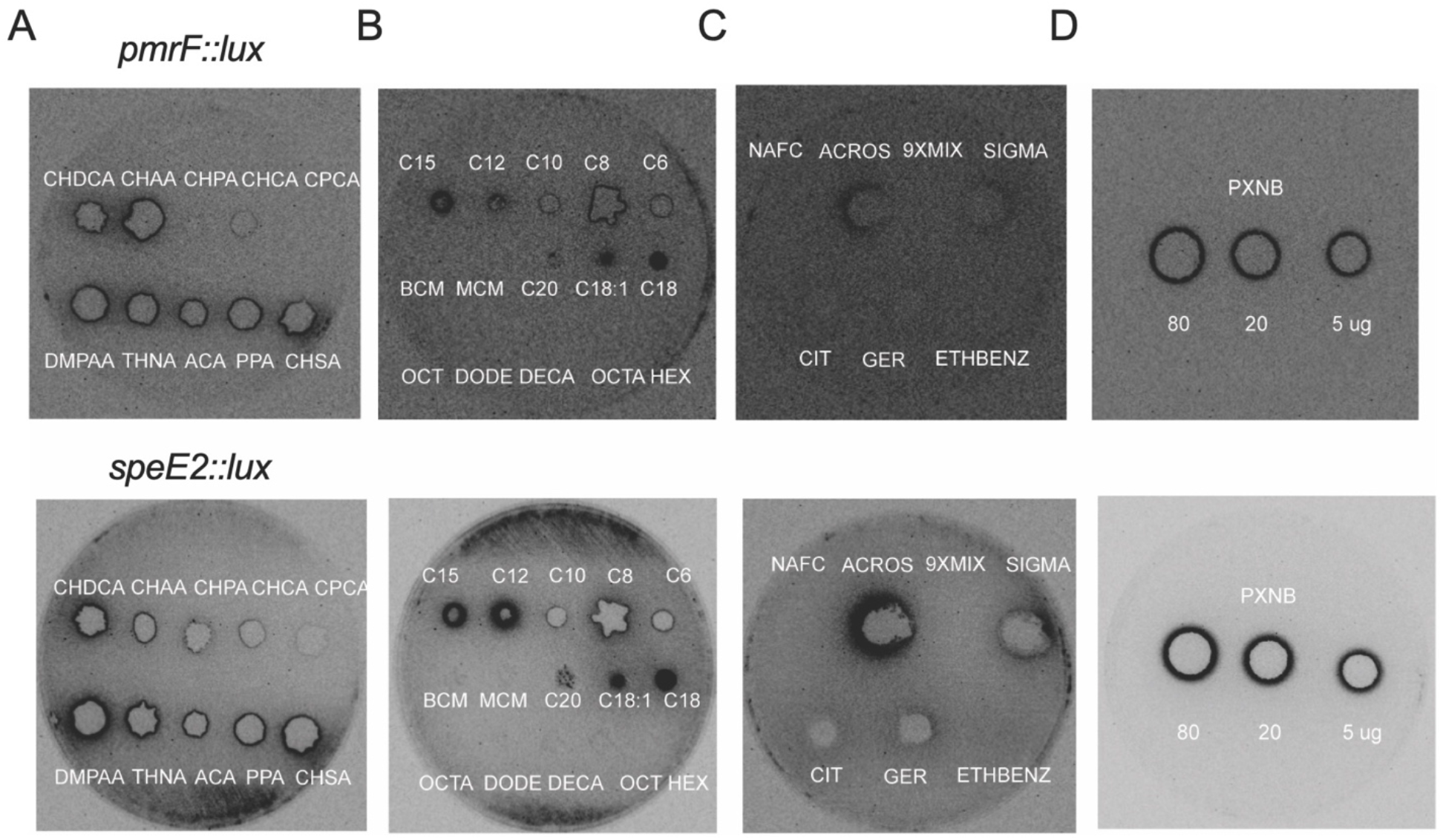
Luminescent rings from naphthenic acids spotted onto a lawn of biosensor culture. Four µL of a 20 mg/ml stock of naphthenic acid compound or mixture was spotted onto a bacterial biosensor lawn. The bioluminescence pattern is imaged after an overnight incubation at 37°C. Images were inverted such that white luminescence patterns are shown as black rings. A) Naphthenic acids B) Fatty acids and alkanes C) NA mixtures, terpenes, ethylbenzene D) Polymyxin B control. See Table 1 for full names. The top and bottom rows of images are from the *pmrF::lux* and *speE2::lux* biosensors, respectively. Exposure times were 180 sec for all naphthenic acids and only 30 sec for polymyxin B.

For all compounds, 80 µg were added and therefore the relative size of the zone of killing can be indicative of the antimicrobial activity. For example, both *P. aeruginosa* transcriptional reporters were very sensitive to naphthenic acids (Fig 3A) and less sensitive to fatty acids (Fig 3B), as observed by the inner ring dimensions for all spotted samples. The strongest fatty acid inducers of outer membrane damage were those ranging in length C12-C18, but C8-C10 fatty acids were more antimicrobial (Fig 3). Of the four NA mixtures tested, the strongest responses were to commercially available NA that are known to contain abundant acyclic fatty acids (28), relative to the cyclic components (Fig 3C). Polymyxin B is a cyclic peptide antimicrobial compound that acts as a positive control. It is an efficient antibiotic with a low MIC (~1 µg/mL) and thus shows a large zone of killing, and a bright ring of luminescence with a short exposure time (Fig 3D). By comparison, 80 µg of the NAFC and 9xNA mixtures were not toxic or detected on agar, likely due to the dilution of membrane-active compounds within the mixtures (Fig 3).

In summary, naphthenic acids and fatty acids induced expression of the biosensors, which indicates the ability to damage the outer membrane (19). Fatty acids were only detected in the solid surface assay. Additionally, the total amount of compound added was greater in the agar plate assay (80 µg) compared to liquid assays (12.5 µg). Therefore, the concentration, solubility or gradual diffusion rates across the agar may have increased bioavailability of the fatty acid compounds.

### pH influences the strength of the biosensor responses to naphthenic acids, likely through influencing the state of protonation

Naphthenic acids are amphipathic compounds (charged and hydrophobic) that can interact with and disrupt the bacterial outer membrane. The pH will affect the protonation state of the carboxylic acid, and therefore we wanted to determine the influence of pH on naphthenic acids and their ability to disrupt the membrane. Upon addition of NA to BM2 liquid media, the pH was reduced from 6.5 to more acid pH values of 6 to 5.5 (Table 2). The *pmrF::lux* biosensor responses to naphthenic acids were compared with and without balancing the pH of growth media. Under neutral pH 6.5 conditions, the biosensor responses to naphthenic acids either did not change, increased, or decreased, depending on the compound (Table 2). There was no consistent biosensor response to a decrease in pH. This suggests that pH can influence the protonation state of naphthenic acids, which can in turn affect the ability to interact with and destabilize membranes. Many naphthenic acids span a range of intrinsic pKa values between 4-5. At biologically relevant pH (6–8), this places many compounds above the pKa values, which produces mostly the anionic state. Membrane disruption is likely affected by hydrophobicity, ring composition, alkyl chain length or branching, and charge. Therefore, not all NA become better membrane disruptors at neutral pH, despite having a higher proportion of anions.

**Table 2.**
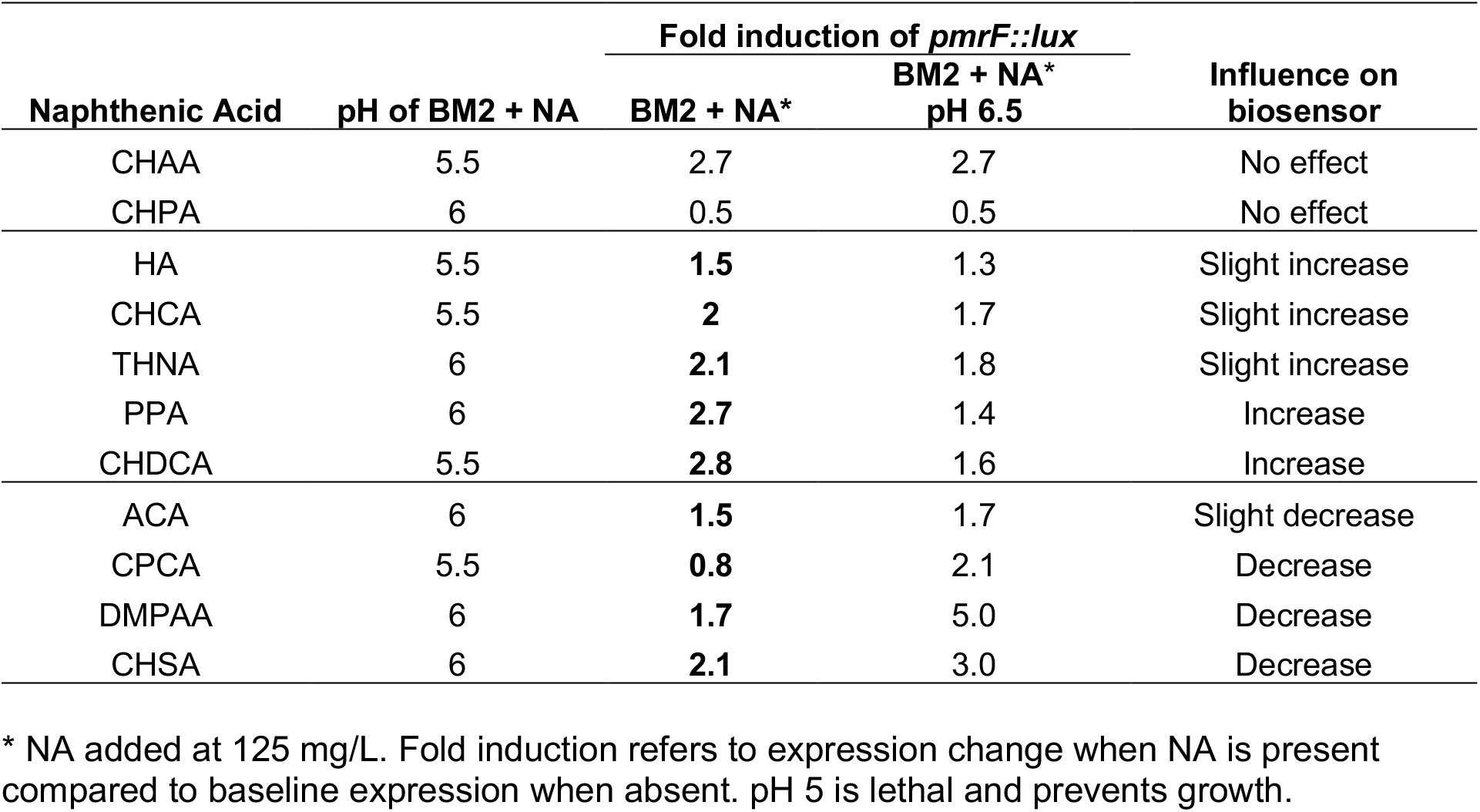
Influence of naphthenic acids on media pH and biosensor responses.

| Naphthenic Acid | pH of BM2 + NA | Fold induction of *pmrF::lux* |  | Influence on biosensor |
| --- | --- | --- | --- | --- |
|  |  | BM2 + NA* | BM2 + NA* pH 6.5 |  |
| CHAA | 5.5 | 2.7 | 2.7 | No effect |
| CHPA | 6 | 0.5 | 0.5 | No effect |
| HA | 5.5 | **1.5** | 1.3 | Slight increase |
| CHCA | 5.5 | **2** | 1.7 | Slight increase |
| THNA | 6 | **2.1** | 1.8 | Slight increase |
| PPA | 6 | **2.7** | 1.4 | Increase |
| CHDCA | 5.5 | **2.8** | 1.6 | Increase |
| ACA | 6 | **1.5** | 1.7 | Slight decrease |
| CPCA | 5.5 | **0.8** | 2.1 | Decrease |
| DMPAA | 6 | **1.7** | 5.0 | Decrease |
| CHSA | 6 | **2.1** | 3.0 | Decrease |

\* NA added at 125 mg/L. Fold induction refers to expression change when NA is present compared to baseline expression when absent. pH 5 is lethal and prevents growth.

### Naphthenic acids disrupt bacterial membranes leading to enhanced lysosome killing or uptake of fluorescent DNA binding dyes

Additional approaches were considered to validate the narcosis activity of naphthenic acids. N-phenyl-1-naphthylamine (NPN) uptake is a conventional method to measure outer membrane disruption, where treatment of a cell with a membrane-disrupting compounds causes a rapid increase in NPN fluorescence (19, 25). NPN is only fluorescent when it partitions in the hydrophobic area of a disrupted membrane (25). We attempted to use the NPN uptake assay as a method to confirm the direct membrane disruption by naphthenic acids. However, when NPN is mixed only with naphthenic acids, alkanes or oleic acid, there is a strong increase in fluorescence in the absence of bacterial cells (Fig 4A). This result suggests the formation of hydrophobic micelles or aggregates, that lead to NPN fluorescence. When NPN is added to cells and then treated with NA, the NPN fluorescence decreased (Fig 4B). This result suggested NPN was sequestered by NA aggregates, and contrasts with the positive control treatment (polymyxin B) that causes an increased NPN fluorescence. Generally, membrane disruption allows greater access of NPN to the membrane (Fig 4B). For these reasons, the NPN uptake is not suitable for the study of hydrophobic compounds such as NA, fatty acids or alkanes, as NPN fluorescence is stimulated without cells.

**Figure 4.**
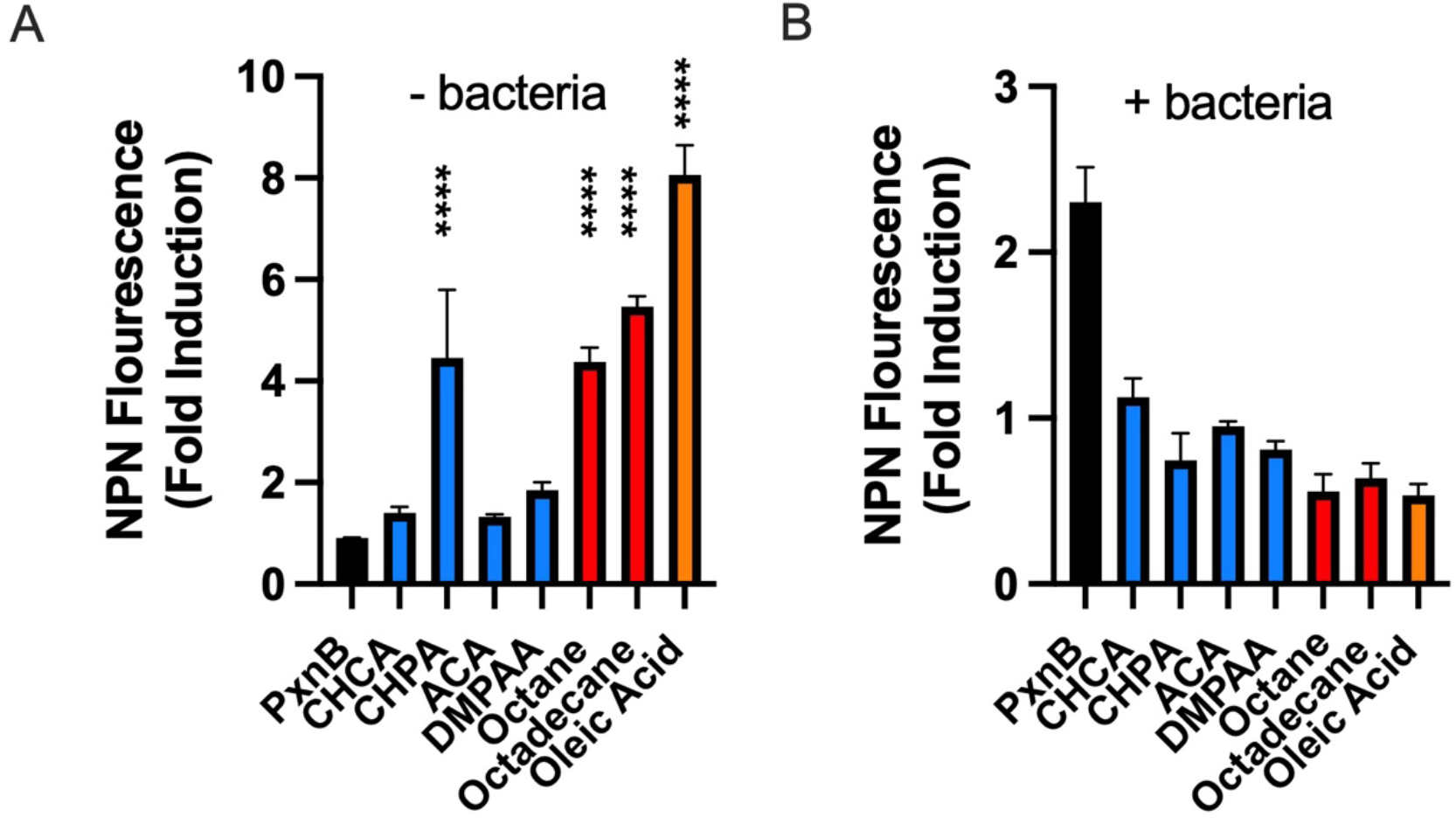
The NPN uptake assay is not a reliable assay for measuring outer membrane disruption by naphthenic acids. **A)** Fold change values of NPN fluorescence increased in the presence of different NA compounds without bacterial cells. **B)** When *pmrF::lux* bacterial cells were added, NA compounds caused a decrease in fluorescence. All compounds were tested at 500 µg/ml, and each value shows an average and standard deviation of three replicates. All experiments were repeated three times. Data were analyzed by one-way ANOVA to test for an overall treatment effect, followed by Dunnett’s multiple-comparisons test to compare each treatment with the control. Significant increases in NPN fluorescence relative to the control is shown with asterisks (**** p < 0.0001).

We therefore used alternative approaches to directly measure the ability of naphthenic acids to disrupt the bacterial membrane. Lysozyme is an enzyme that degrades the peptidoglycan cell wall located in the periplasm, which is normally protected by the outer membrane. In the lysozyme-assisted killing assay, compounds are tested for permeabilizing the outer membrane barrier, which permits lysozyme to enter and degrade the cell wall, leading to lysis and killing (11, 27). Figure 5A depicts the control condition of lysozyme alone that does not kill *P. aeruginosa* or reduce the optical density, but the addition of both polymyxin B and lysozyme leads to killing and drop in OD_600_. Increasing concentrations of polymyxin B and lysosome produced increasing levels of bacterial killing (Fig 5B). The individual NA compound cyclohexane acetic acid (CHAA), and the simple 9xNA mixture were also shown to increase lysozyme-assisted killing (Fig 5B). This supports the interpretation that naphthenic acids permeabilize the outer membrane and allows lysozyme to enter the periplasm and degrade the peptidoglycan cell wall. In general, naphthenic acids promoted weaker killing than the OM targeting antimicrobial peptide polymyxin B (Fig 5), which is consistent with naphthenic acids being weak or moderate inducers of the *pmrF::lux* biosensor, relative to polymyxin B (Fig 3) and other OM targeting compounds (19).

**Figure 5.**
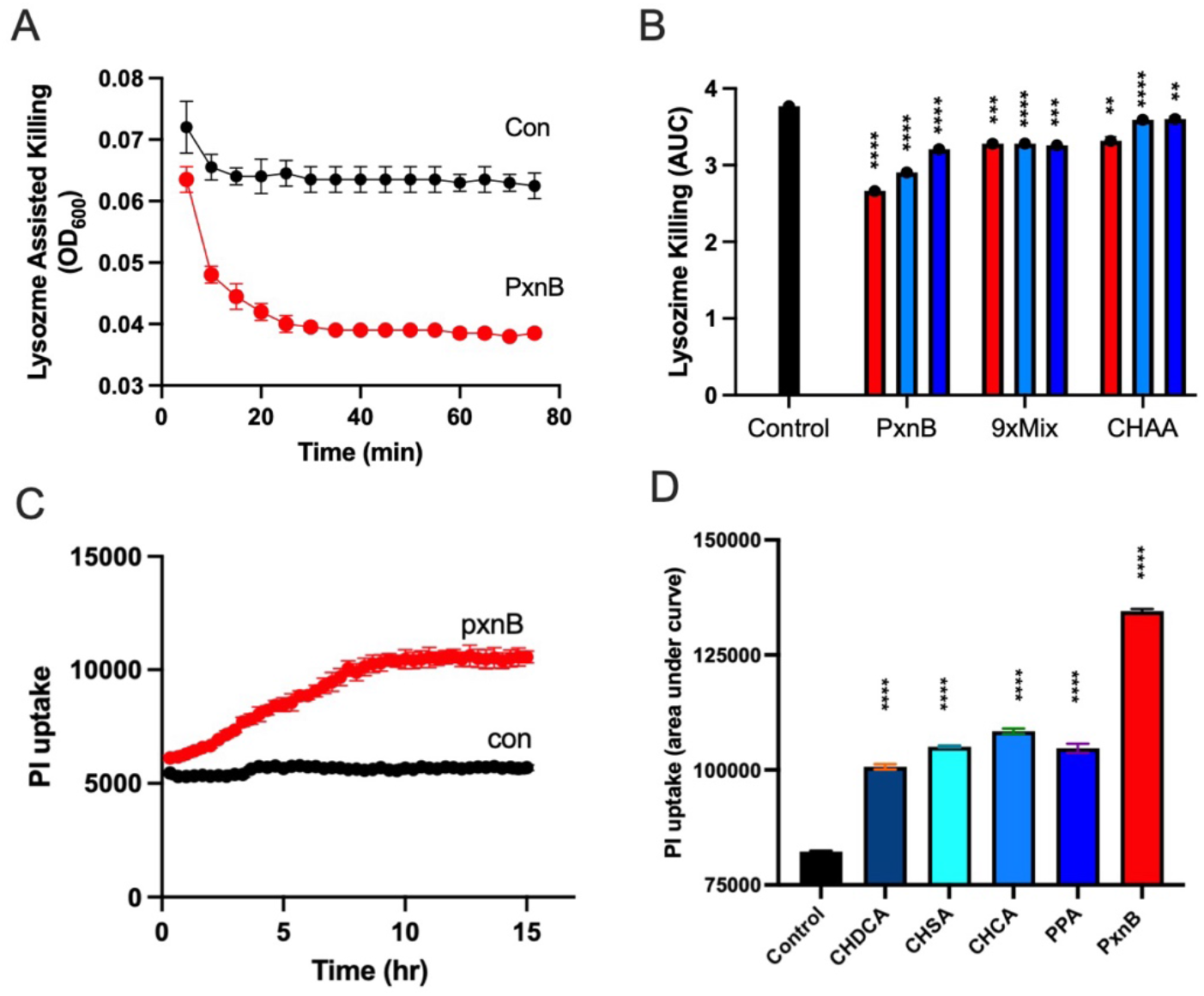
Naphthenic acids disrupt the integrity of bacterial cell membranes. The lysozyme killing assay (**A, B**) or the propidium iodide uptake assay (**C, D**) were used to assess the membrane integrity after naphthenic acids exposure. **A)** Lysis (OD_600_) was measured after treating bacterial cells with lysozyme alone (black) or lysozyme with 10 µg/mL polymyxin B (red). Measurements were taken every 5 minutes for a total of 75 minutes. **B)** Area under the curve (AUC) of the lysis measurements (OD_600_) during the 75-min assay was calculated for each compound and concentration tested with lysozyme. The 9xNA mixture and CHAA were both tested at 1 mg/mL (red), 0.5 mg/mL (light blue), and 0.25 mg/mL (dark blue). PxnB was tested at 10 µg/mL (red), 5 µg/mL (light blue), and 2.5 µg/mL (dark blue). Values shown are an average and standard deviation of two replicates. **C)** Red fluorescence due to PI uptake after treating the bacterial cells with polymyxin B. Measurements taken every 20 minutes for 15 hours. **D)** AUC of the red PI fluorescence during the 15-hour assay were calculated for each compound. PxnB was tested at 1 µg/mL and each NA compound was tested at 125 mg/L. Values shown are an average and standard deviation of three replicates. Data were analyzed by one-way ANOVA to test for an overall treatment effect, followed by Dunnett’s multiple-comparisons test to compare each treatment with the control. Significant membrane disruption relative to the control is shown with asterisks (** p < 0.01, *** p < 0.001, **** p < 0.0001). All experiments were repeated three times.

To determine if naphthenic acids could disrupt both the outer and inner membranes in *P, aeruginosa*, we monitored the uptake of the DNA binding dye propidium iodide. PI is a membrane-impermeable compound that does not enter the cell in undisturbed cells, but polymyxin B causes a significant increase in the uptake of PI (Fig 5C, 5D). Naphthenic acids were also shown to promote significant levels of PI uptake in *P. aeruginosa*, albeit at lower levels than polymyxin B (Fig 5D), consistent with the effects of NA treatment of the bacterium *R. aertherivorans* (10) or in onion epidermal cells (9). Taken together, these assays provide further evidence that naphthenic acids interact with and disrupt bacterial membranes, consistent with their proposed narcosis mechanism of action.

## Discussion

Naphthenic acids (NAs) are widely hypothesized to exert toxicity through narcosis, a mechanism in which amphipathic compounds partition into and disrupt biological membrane function. Despite this long-standing model, relatively few assays directly measure bacterial membrane disruption by NA mixtures or individual NA constituents. Here, we demonstrate that NAs can be detected as membrane-active, narcosis stressors using a whole-cell biosensor platform based on the *Pseudomonas aeruginosa* envelope defense pathways. Two independent transcriptional reporters, *pmrF::lux* and *speE2::lux*, were induced by a broad panel of individual model NA compounds, NA mixtures, and OSPW-derived extracts. Reporter induction was supported by functional assays demonstrating envelope compromise, including lysozyme-assisted killing and propidium iodide uptake. In addition to direction partitioning into hydrophobic membranes, naphthenic acids can associate with Mg^2+^ and Ca^2+^ to form metal naphthenates that are found at the interface of oil and water (29). This activity may potentially disrupt cation interactions on the bacterial surface, similar to cation chelators (11, 17). Together, these findings support the narcosis hypothesis and establish a novel, high throughput approach to detect NA membrane disruption (Fig 6).

**Figure 6.**
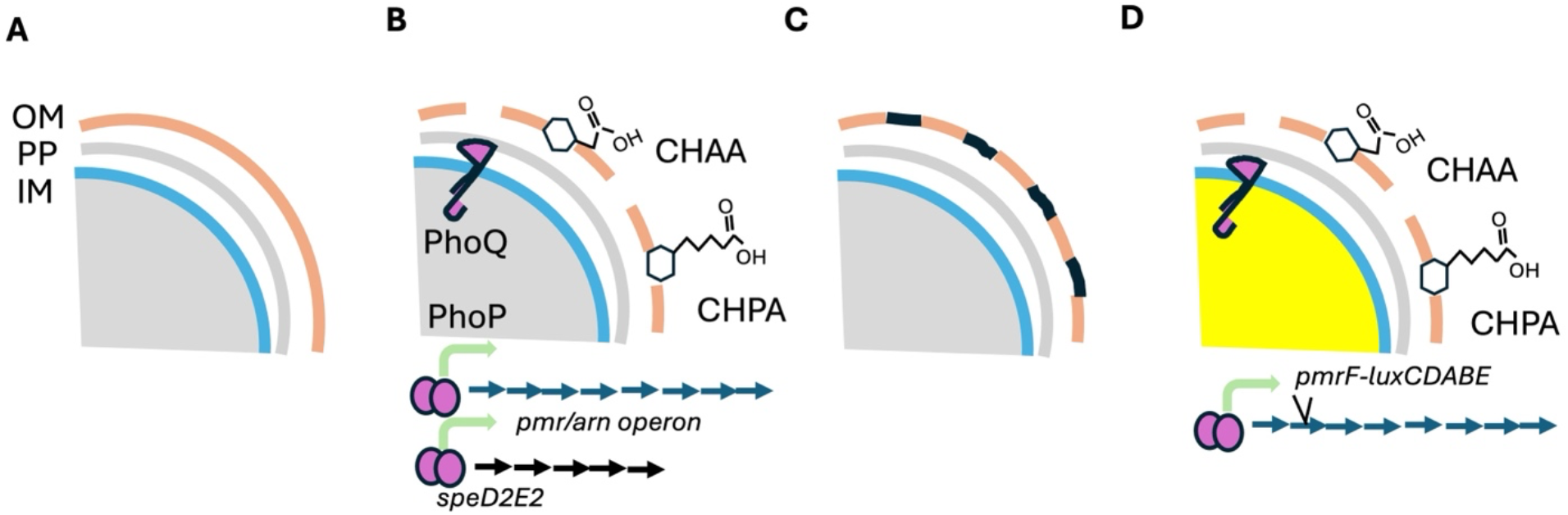
Narcosis effects on the bacterial outer membrane leads to the expression of protective outer membrane repair processes. **A.** In normal conditions, the cell envelope consisting of the outer membrane (OM), the periplasm (PP) and inner membrane (IM) is intact. **B**. Naphthenic acids are hydrophobic and can partition into and disrupt the OM integrity or can potentially disrupt surface stabilizing cations. This damage is sensed by various two-component regulatory systems (eg PhoPQ, PmrAB), which include an inner membrane sensor protein. Upon activation, the sensor phosphorylates its cognate response regulator, which now upregulates the expression of target genes. In this case, the *pmr/arn* operon and *speD2E2* operons are induced. **C**. The outer membrane is repaired by covalent modification of aminoarabinose to the phosphates of LPS, and the presence of spermidine, a polycation that replaces natural divalent cations such as Mg^2+^. Membranes expressing these modifications are tolerant to future outer membrane narcosis. **D**. A *luxCDABE* cassette is inserted into the *pmrF*/*arnC* gene, which produces luminescence upon OM disruption by NA.

The advantages of this narcosis biosensor assay include a rapid and simple assay, real-time gene expression measurements with no added substrates, luminescence responses specifically to membrane damage rather than generic toxicity (eg Microtox), sensitive detection of membrane damage at sub-lethal concentrations of NA, high throughput capacity to screen many compounds or water samples containing NA under varying parameters (concentrations, solvents, pH), semi-quantitative detection of membrane damage based on gene expression values (CPS), a wide dynamic range to compare strong and weak activity, and lastly is suitable for hydrophobic compounds that prevent use of the conventional NPN uptake assay. The limitations of this narcosis bioassay include possible false negatives when testing volatile hydrocarbons, broad narcosis detection rather than specific naphthenic acid detection, and the method requires further development as a direct indicator of toxicity of OSPW.

We previously reported NA-responsive regulatory biosensors (e.g., AtuR/atuA and a MarR-based system) to report on total NA concentration. Each biosensor detects a specific subset of bioavailable NA compounds once inside cells, through ligand-responsive transcriptional repressors (21). In contrast, the *pmr/spd* biosensors respond to surface membrane disruption, and these genes are controlled by numerous two-component regulatory systems (TCS) consisting of membrane-bound sensors and cytoplasmic regulators, including PhoPQ, PmrAB, and potentially CprRS or ParRS (Fig 5). PhoPQ and PmrAB are important for the response to limiting Mg^2+^ or acid stress (14, 18, 30), while CprRS and ParRS were described as important for the sensing of antimicrobial peptides directly (31, 32). All conditions that lead to upregulation of the *pmr* and *spd* operons may in fact be membrane stress conditions. This highlights the complexity of bacterial NA detection mechanisms, where complex NA mixtures may contain degradable compounds, intracellular toxic compounds requiring efflux, and narcosis causing compounds that target the membrane. Future work should resolve how these regulatory TCS discriminate among different classes of membrane permeabilizing agents.

The narcosis biosensor approach detects bacterial membrane disruption and aligns conceptually with biomimetic extraction using solid-phase microextraction (BE-SPME), which recovers potentially narcosis causing NA compounds from OSPW that partition into synthetic, hydrophobic membranes (8, 33). In future work, the narcosis biosensors may be developed to assess OSPW toxicity by comparing the biosensor and BE-SPME methods and correlating their output to toxicity endpoints using various animal models. In summary, these results provide direct evidence that naphthenic acids disrupt bacterial membranes and offers a sensitive and scalable assay platform for detecting narcosis activity by naphthenic acids.

## Funding

This work was supported from a Genome Canada Large-Scale Applied Research Project (LSARP) grant (#18207), in partnership with Genome Alberta and Genome Quebec. Co-funding was provided by the Government of Alberta through an Alberta Innovates grant and Jobs, Economy and Innovation funding.

## Notes

### Competing Interest Statement

The authors have declared no competing interest.

